# A Signal Demodulation-based Method for the Early Detection of Cheyne-Stokes Respiration

**DOI:** 10.1101/723502

**Authors:** Pauline Guyot, El-Hadi Djermoune, Bruno Chenuel, Thierry Bastogne

**Affiliations:** CRAN UMR 7039, Université de Lorraine, CNRS, Vandœuvre-lès-Nancy, France; CYBERnano, 193 avenue Paul Muller, 54602 Villers-lès-Nancy, France; EA 3450 DevAH, Université de Lorraine, Vandœuvre-lès-Nancy, France; INRIA, BIGS, Vandœuvre-lès-Nancy, France

## Abstract

Cheyne-Stokes respiration (CSR) is a sleep-disordered breathing characterized by recurrent central apneas alternating with hyperventilation exhibiting a crescendo-decrescendo pattern of tidal volume. This respiration is reported in patients with heart failure, stroke or damage in respiratory centers. It increases mortality for patients with severe heart failure as it has adverse impacts on the cardiac function. Early stage of CSR, also called periodic breathing, is often undiagnosed as it only provokes hypopneas instead of apneas, which are much more difficult to detect. This paper demonstrates the proof of concept of a new method devoted to the early detection of CSR. The proposed approach relies on a signal demodulation technique applied to ventilation signals measured on 15 patients with chronic heart failure whose respiration goes from normal to severe CSR. Based on a modulation index and its instantaneous frequency, oscillation zones are detected and classified into three categories: CSR, periodic breathing and no abnormal pattern. The modulation index is used as an efficient biomarker to quantify the severity of the pathology for each patient. Results show high correlation with experts’ annotations with sensitivity and specificity values of 87.1% and 89.8% respectively. A final decision leads to a classification which is confirmed by the experts’ conclusions.

## 1 Introduction

Cheyne-Stokes respiration (CSR) is a type of sleep-disordered respiration characterized by a crescendo-decrescendo pattern of ventilation, alternating hyperventilation and central hypopneas/apneas. CSR is mainly prevalent in patients with severe heart failure (left ventricular ejection fraction less than 30%) and can be associated with a worse prognosis [1, 2]; but it can also be found in patients with history of stroke, exposure to high altitude or damages in respiratory centers. Previous investigations have shown that Central Sleep Apnea (CSA) associated to CSR is a strong independent marker of mortality in patients with heart failure [1], and there is an intense need for developing better diagnostic and prognostic tools in order to generate personalized medicine with new and effective treatments [3].

The home respiratory polygraphy (HRP) is probably the most used ambulatory test to identify sleep disorders such as CSR. HRP requires a portable device to record multiple physiological parameters throughout the night, such as blood oxygen saturation, heart rate, airflow, thoracic effort, abdominal effort and body position. HRP is often used as an alternative to an in-hospital test where the overnight multi-channel polysomnography (PSG) [4] is recognized as the reference method to identify patients with periodic breathing (PB) preceding CSR and apnea. This multiparametric test monitors many other body activities such as brain activity (electroencephalogram), eye movements (electrooculogram), muscle activity or skeletal muscle activation (electromyogram) and heart rhythm (electrocardiogram) during sleep. Unfortunately, its average cost is about five times more expensive than HRP.

Several clinical studies have assessed and compared HRP and PSG in order to diagnose CSR. In 2004, a clinical trial applied to 75 patients showed that HRP had a high sensitivity and specificity for the diagnosis of sleep-disordered breathing associated with heart failure [5]. In [6], authors carried out another clinical study over about 350 patients and confirmed that HRP was an efficient alternative to polysomnography in patients with suspected sleep apnoea-hypopnoea syndromes. Nevertheless, Alonso-Álvarez *et al.* emphasized in [7] that HRP was indeed a reliable approach for the diagnosis of obstructive sleep apnea but more research is required for the diagnosis of mild syndromes. In another study published in 2014, Tan *et al.* [8] also revealed that apnea-hypopnea index (AHI), the standard measure to evaluate CSR or periodic breathing, is underestimated in HRP and that the disparity of HRP and PSG indexes can significantly affect clinical management decisions, particularly in children with mild and moderate obstructive sleep apnea. Those recent studies emphasize the difficulty to detect early patterns of sleep disorders like CSR.

The recurring problem is to detect significant amplitude oscillations among the respiratory signals. Some methods have been proposed to quantify the amplitude of the oscillations. For example, a spectral decomposition algorithm of the instantaneous minute ventilation is proposed in [17–19] where periodic breathing has to be previously detected to be quantified. A method based on a standard amplitude demodulation scheme based on filters is presented in [16]. Those two methods can be noise-sensitive and only bring information on the amplitude of the modulation but does not specify any pattern characteristics such as the instantaneous frequency of the oscillation, thus cannot confirm a CSR pattern.

The objective of this paper is to propose a novel computational method able to better detect and classify early patterns of CSR in respiratory signals in order to improve an early diagnosis and to propose an index to quantify the severity of the pathology. The proposed algorithm does not need to previously detect periodic breathing to quantify it. The whole respiratory signal is processed and the algorithm is able to specify zones of interest. Our contribution relies on a signal amplitude modulation technique which is well suited to the crescendo-decrescendo pattern of CSR. The estimated modulation index is used as a biomarker to estimate the CSR stage. A panel of 15 patients with chronic heart failure was used to demonstrate the proof of concept. To assess the performances of the local detection and final classification, the results obtained by the new method were compared to those given by eAMI [16] and the opinion of CSR experts.

The remainder of the paper organized as follows. Section 2 describes the respiration model based on amplitude modulation for the estimation of our indices. Then, the details of the proposed computational method are presented in Section 3. The results obtained on a panel of fifteen patients are presented in Section 4 and compared with sleep experts. They are discussed in Section 5. Finally, conclusions are drawn in Section 6.

## 2 Amplitude modulation-based model of Cheyne-Stokes Respiration

Amplitude modulation is mainly used in radio transmission for broadcasting and communication. Two signals are used to create a modulated signal: the carrier wave, which is a high frequency signal and the information-bearing modulation signal of lower frequency. A modulated signal is obtained by varying the amplitude of the carrier wave with the modulation signal. In the case of CSR, the ventilation can be modeled as follows (see Figure 1 for a graphical illustration):

**Fig 1.**
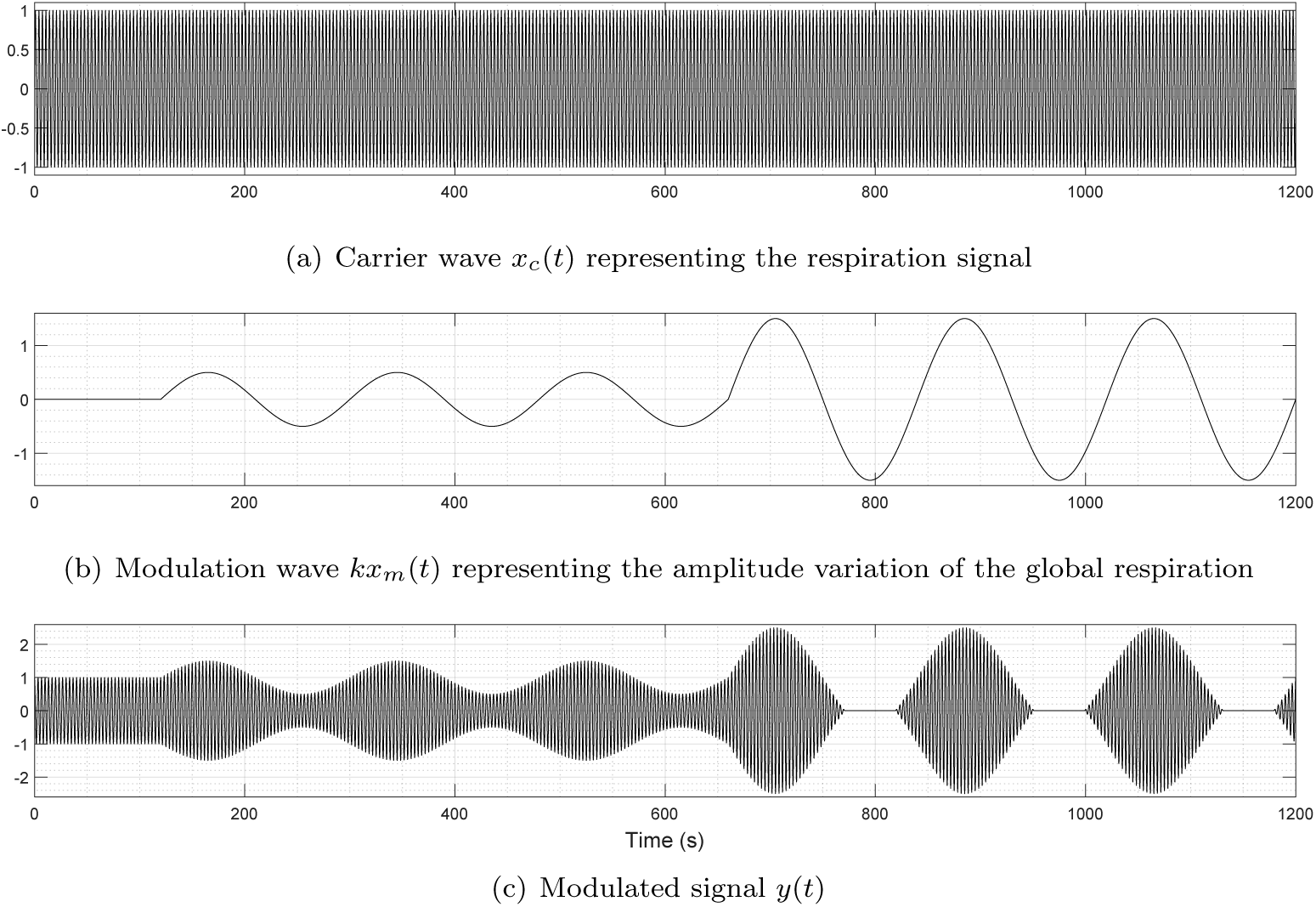
Modeling of Cheyne-Stokes Respiration *via* amplitude modulation. The modulated signal in (c) shows a normal respiration for the first two minutes, then an early stage of Cheyne-Stokes Respiration (*h* = 0.5) and a severe form of Cheyne-Stokes respiration (*h* = 1.5) at *t* = 660 s = 9 min.

- the carrier wave represents the respiration signal and is considered as a sinusoidal signal *x*_*c*_(*t*) whose frequency *f*_*c*_ goes from 0.25 Hz to 0.33 Hz in the case of CSR (from 15 to 20 respirations per minute for adults):

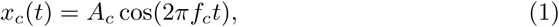

where *t* denotes the time variable and *A*_*c*_ the carrier amplitude;
- the modulation signal which stands for the enveloppe of the respiration, is either constant for a normal respiration or oscillating for a CSR pattern. Similarly, it is also assumed to be a sinusoidal signal *x*_*m*_(*t*) whose frequency goes from 8 mHz to 30 mHz (a cycle of CSR typically lasts from 30 s to 2 min):

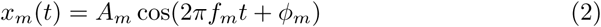

where *A*_*m*_ is the modulation amplitude and *ϕ*_*m*_ is the phase;
- the modulated signal can be expressed as:

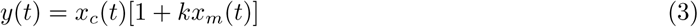

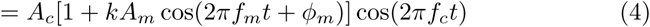

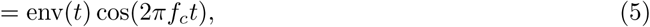

where *k* ∈ ℝ is a constant and *h* = *kA*_*m*_ is the modulation index. The enveloppe signal is defined as follows:

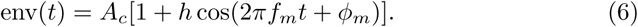

The modulation index *h ≥* 0 is a key parameter of the amplitude modulation. It varies between 0 and 1 and graduates the amplitude level of the periodic breathing as indicated in Figure 1. Over-modulation (*h >* 1) creates a distortion of the signal, but this case is not considered here because it cannot happen for respiration. However, apnea can occur and the modulated signal is modified to:

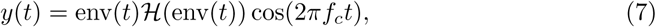

where ℋ (*t*) is the Heaviside function defined by ℋ (*t*) = 1 for *t >* 0 and ℋ (*t*) = 0 otherwise. Note that the function ℋ (·) in (7) is effective only when *h >* 1, otherwise it is equal to 1. When *h >* 1, the duration *δ* of the apnea period can be computed from *h* and *f*_*m*_:

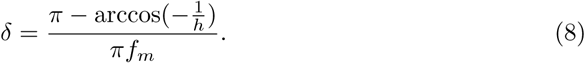

Figure 1 shows an apnea zone around *t* = 800 s corresponding to *h* = 1.5.

## 3 Proposed algorithm

The proposed computational method can be decomposed into four successive steps: (i) computation of the envelope of the ventilation signal; (ii) estimation of the modulation index and its instantaneous frequency; (iii) detection of potential CSR or periodic breathing zones; and (iv) final classification for each patient in three different categories: CSR, periodic breathing or non-CSR. These steps are thoroughly described in the next section.

### 3.1 Envelope computation

This part of the method is composed of two stages: (1) detection of breathing cycles in the ventilation signal and (2) reconstruction of the envelope.

#### Detection of breathing cycles

Change point analysis (CPA) [20] is used to detect breath-by-breath respiration from the ventilation signal. Let us denote by **x** ∈ ℝ^*N*^ the respiration signal to be analyzed. We assume that some statistical properties of **x** change abruptly at instants *t*_1_, *…, t*_*K*_, called change points. In CPA methods, the aim is to estimate the segmentation 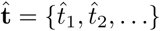 through the minimization of a cost function *C* which represents the sum of squared residuals. When the number of changes *K* is unknown, a penalty term (regularization) is added to the residual error. The approach tend to minimize:

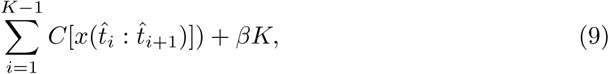

with *β* > 0 is a tuning parameter that controls the number of change points K [23]. In our case, a change point represents a peak or a trough in the signal (inspiration and expiration events) and the statistical properties used are slope and mean. Once all change points are detected, slope is computed for all sections and those lower than a threshold (experimentally set to 10^−3^) are discarded and considered as noise. Finally, only peaks whose section duration is greater than one second are conserved (biological prior knowledge: the respiratory rate is between 15 and 20 cycles per minute). An example of segmentation is given in Figure 2.

**Fig 2.**
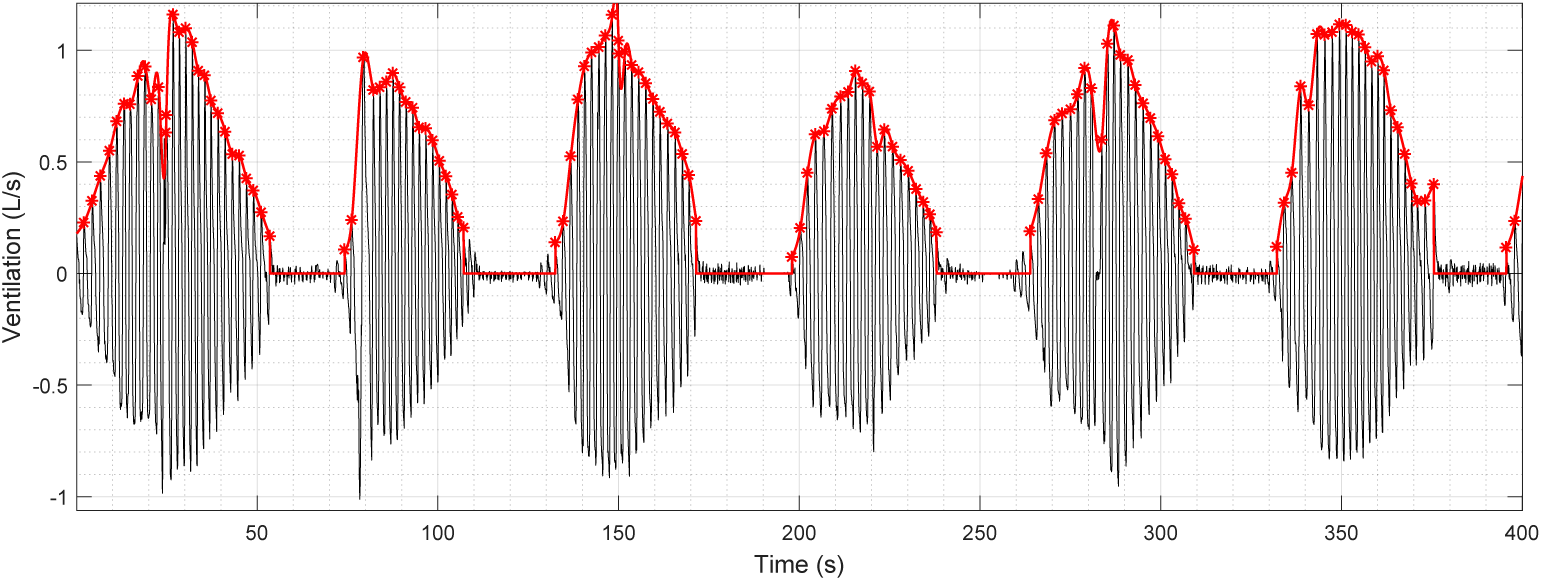
Reconstruction of the ventilation envelope for a patient with severe Cheyne-Stokes respiration. Results of change point detection for breathing cycles are plotted with red stars and the envelope of the ventilation signal with the red line.

#### Reconstruction of the envelope

Interruption of ventilation is detected if the time difference between two breaths is greater than three times the median of the distances between peaks. In this case, the envelope is set to zero until the next breath (see also Figure2). Finally, the signal is linearly interpolated and then evenly resampled.

### 3.2 Parameter estimation of the CSR model

Once the envelope of the ventilation signal is extracted, the goal is to estimate the parameters *A*_*c*_, *f*_*m*_, *ϕ*_*m*_ and *h* of the CSR envelope model presented in (6). As the envelope is modeled as a sinusoidal process, we used a subspace-based method called Matrix Pencil [21, 22]. First, let us express the envelope as a weighted sum of complex exponentials:

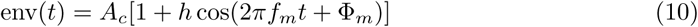

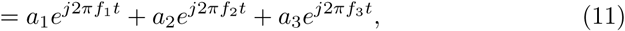

with frequencies *f*_1_ = 0, *f*_2_ = *f*_*m*_, *f*_3_ = −*f*_*m*_ and complex amplitudes *a*_1_ = *A*_*c*_, 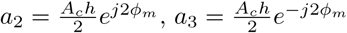, where 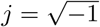. These parameters are then estimated by the Matrix Pencil method over a sliding window. The window size *t*_w_ has to be small enough for the stationarity assumption (the sinusoidal model with locally constant parameters) to hold. Here it is set to 2 minutes and no difference was found for *t*_w_ ∈ [2, 4]. The overlapping ratio *ρ*_w_ between two successive windows is set to 80%.

### 3.3 Detection of CSR zones

According the value of *ĥ = 2*|*â*_2_|*/â*_1_ and 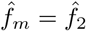 (the hat symbol indicates estimated quantities), a decision is made to decide whether the envelope is constant or oscillating. Through ROC analysis using experts annotations, a threshold of *h*_0_ = 0.12 was used to detect a modulation of breathing sufficiently present to be pathological. In parallel, 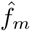 has to belong to the interval [8, 30] mHz in which Cheyne-Stokes pattern is typically pathological. If both *ĥ* and 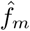 are classified as pathological for at least 1 minute, then a zone of CSR pattern is detected and the value of *ĥ* specifies the severity of the pathology. The one-minute window decision is used to avoid artifacts triggered by short false positives.

### 3.4 Severity classification of the CSR pathology

Based on the American Academy of Sleep Medicine (AASM) recommandations [24], a final classification rule can be applied to each patient to assign a diagnosis:

- if the duration of breathing oscillation is longer than 10 minutes with *ĥ* greater than 1 (at least 5 cycles) with minimum one episode lasting at least 6 minutes (3 consecutive cycles) then the patient is classified as CSR-CSA (severe CSR pattern with apneas);
- if it is longer than 10 minutes but *ĥ* is less than 1 with minimum one episode lasting at least 6 minutes (3 consecutive cycles) then the patient exhibits an early stage of CSR and the value of *ĥ* can be interpreted as an indicator of severity of the pathology;
- if it is shorter than 10 minutes with no episode lasting more than 3 consecutive cycles, the patient is classified as non-CSR.

## 4 Database and results

### 4.1 Study design

This study is a retrospective analysis of data, which included adult patients referred to a sleep laboratory (University Hospital CHRU Nancy) for evaluation of suspected sleep disordered breathing. The study was approved by the Local Ethics Committee of the University Hospital of Nancy and informed consent was obtained from all subjects before they commenced participation. It involves a group of fifteen patients all presenting severe heart failure (LVEF^1^<30%). Patient characteristics are listed in Table 1. Subjects were seated comfortably on a chair in a quiet room, in a condition of relaxed wakefulness for about 30 minutes of recording. They breathed room air through a low-dead-space face mask (Hans Rudolph mask, 7400 oro-nasal series, small or medium size, Hans Rudolph, Kansas City, KS) connected to a pneumotachograph (MediGraphics Prevent pneumotachograph, Medical Graphics, St. Paul, MN). Inspiratory and expiratory flows were measured, and the respiratory gas was continuously sampled from the pneumotachograph for the measurement of expired CO_2_ and O_2_ partial pressure. Oxygen and CO_2_ concentrations were determined by rapidly responding O_2_ and CO_2_ analyzers (Datex analyzers, Medical Graphics, St. Paul, MN). Respiratory flow, PO_2_ and PCO_2_ were digitized at 200 Hz for breath-by-breath calculation of expiration and pulmonary gas exchange. Oxygen saturation, thoracic belt respiration and blood pressure were also simultaneously recorded. Sleep experts were asked to classify each minute of the ventilation signal, based only on visual inspection, into three categories: (1) CSR or PB, (2) No abnormal pattern and (3) Erratic breathing possibly PB. As a second task, experts had to establish diagnosis following international guidelines using all available signals. Four patients had severe CSR, one patient exhibited a periodic breathing preceding CSR-CSA and ten patients were classified as non-CSR breathing. Among the non-CSR class, three patients were marked with a suspicion of periodic breathing typically preceding CSR-CSA but the experts were not able to confirm this diagnosis on the basis of the available signals. No patient was on opioids.

**Table 1.** Clinical characteristics of a group of 15 patients with severe heart failure.

| | Units | Median | Mean ± SD |
| --- | --- | --- | --- |
| Age | years | 58 | $58 \pm 9.3$ |
| Height | cm | 173 | $174.4 \pm 5.3$ |
| Weight | kg | 81.5 | $81.75 \pm 11.39$ |
| BMI | kg/m<sup>2</sup> | 26.6 | $26.8 \pm 4.12$ |
| LVEF | % | 25 | $24.77 \pm 5.95$ |
| pH | | 7.47 | $7.75 \pm 0.03$ |
| SaO<sub>2</sub> | % | 96.3 | $95.65 \pm 1.98$ |
| BP systolic | mmHg | 120 | $116 \pm 13.52$ |
| BP diastolic | mmHg | 60 | $64.66 \pm 8.33$ |
| Dyslipidemia | % |  | 73.3 |
| Myocardial infarction | % |  | 53.3 |
| Hypertension | % |  | 40 |
| Diabetes | % |  | 20 |
| Hyperthyroidism | % |  | 13.3 |
| Stroke | % |  | 6.66 |

BMI: Body Mass Index. LVEF: Left Ventricle Ejection Fraction. BP: Blood Pressure. Sex: all male.

### 4.2 Diagnostic criteria for Cheyne-Stokes Respiration

Cheyne-Stokes respiration was defined by the presence of the classical pattern of waxing/waning in the tidal volume associated with central hypopneas/apneas. Central hypopnea was defined as a reduction of tidal volume of at least 30% along with a drop of 3% in oxygen saturation – if no flow limitation or obstructive apnea is observed. Central apnea was defined as a cessation of tidal volume for at least 10 seconds without any respiratory efforts.

### 4.3 Competing methods

The eAMI method [16] is the closest proposition to ours in the literature: both methods rely on amplitude modulation but differ in the employed techniques and the computed indexes. It has been completely implemented in MATLAB R2018a for comparison with the proposed method. Figure 3 illustrates the scheme used in [16] for the computation of the eAMI index with the value of each parameter. Using a filter bank, a modulation index is estimated; it indicates an apnea when its value is close to one and can take negative values in the absence of periodic breathing. It requires five main steps and a set of several parameters including the cut-off frequency of each filter, and only a part of them is specified in [16]. In comparison, our method requires three steps that may require more computation resources but with more easily determinable parameters: the set of cut-off frequencies would depend on the nature of the signals while our only critical parameters are the window size and overlapping ratio which are related to the cut-off frequency of the last low-pass filter in eAMI algorithm.

**Fig 3.**
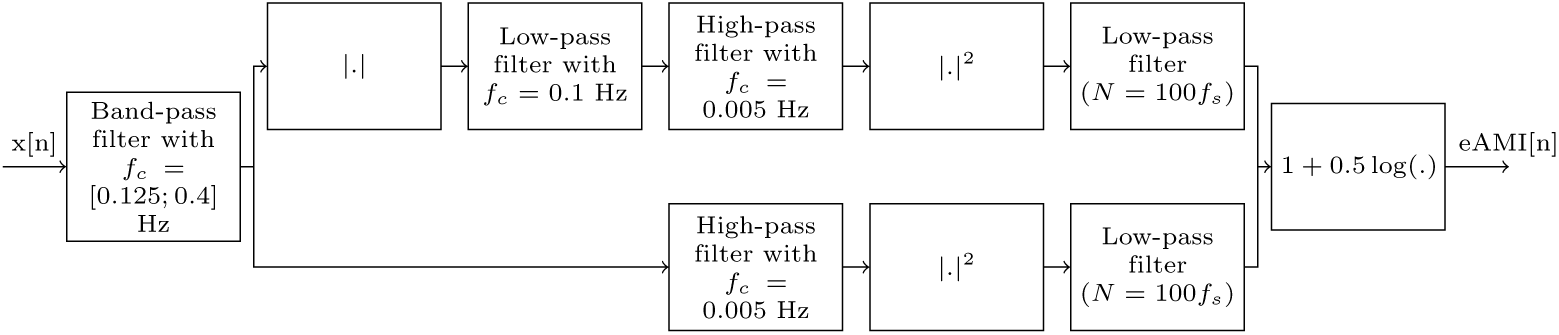
Scheme, taken from [16], used to compute eAMI index. The input x stands for the respiratory signal and the output eAMI for the computed index.

Both eAMI and the proposed method involve a threshold parameter to classify respiration intervals using the computed indexes. They are determined through ROC analysis using experts’ annotations on the first two classes CSR/PB or no abnormal pattern. The parameters selected to assess the performance of zones detection are:

- Sensitivity (Se): 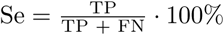
- Specificity (Sp): 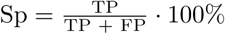

where TP, FP and FN stand for the number of true positives, false positives and false negatives, repectively. A true positive correspond to a minute correctly detected by the algorithms as an oscillation zone. A false positive correspond to a minute detected part of zone meanwhile it is not labeled the same by experts. A false negative correspond to a minute undetected by the algorithms. Finally, the global performance is assessed by computing a confusion matrix for both methods.

### 4.4 One-minute estimation results

Figures 4 and 5 present the estimation results for two patients with different profiles. Each figure shows the raw ventilation signal and the values of *ĥ* and 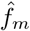 estimated over the sliding window. In Figure 4, a patient with severe CSR exhibits a modulation index above 1, which highlights the presence of apnea and its modulation frequency belongs to the pathological interval of CSR. Combined together, *ĥ* and 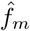 clearly indicate that the patient’s ventilation oscillates with apnea at a pathological frequency during all recording: a zone is detected and classified as CSR. eAMI method also detects correctly CSR even if it stagnates around 0.5 and does not reach the apnea threshold of 1. In Figure 5, *ĥ* is above the oscillation threshold and 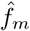 is included in the pathological interval for parts of the recording. The two parameters enable to conclude that the patient’s ventilation shows a modulation in amplitude at a pathological frequency and can be classified as periodic breathing typically preceding CSR. When the envelope of the ventilation signal remains constant, *ĥ* goes under the oscillation threshold: the patient’s ventilation shows no pathological modulation. Concerning eAMI index, it correctly detects the normal episode of respiration but does not perform well for the modulation from *t* = 1000 s. Using all patients, our method achieves a specificity of 89.8% and a sensitivity of 87.11% when compared to experts for the classification on the minute ventilation for the two first classes (CSR/PB or no CSR/PB). For comparison, eAMI achieves a specificity of 78.41% and a sensitivity of 76.44%. Note that, as for the proposed method, the threshold used for eAMI classification is also determined through ROC analysis.

**Fig 4.**
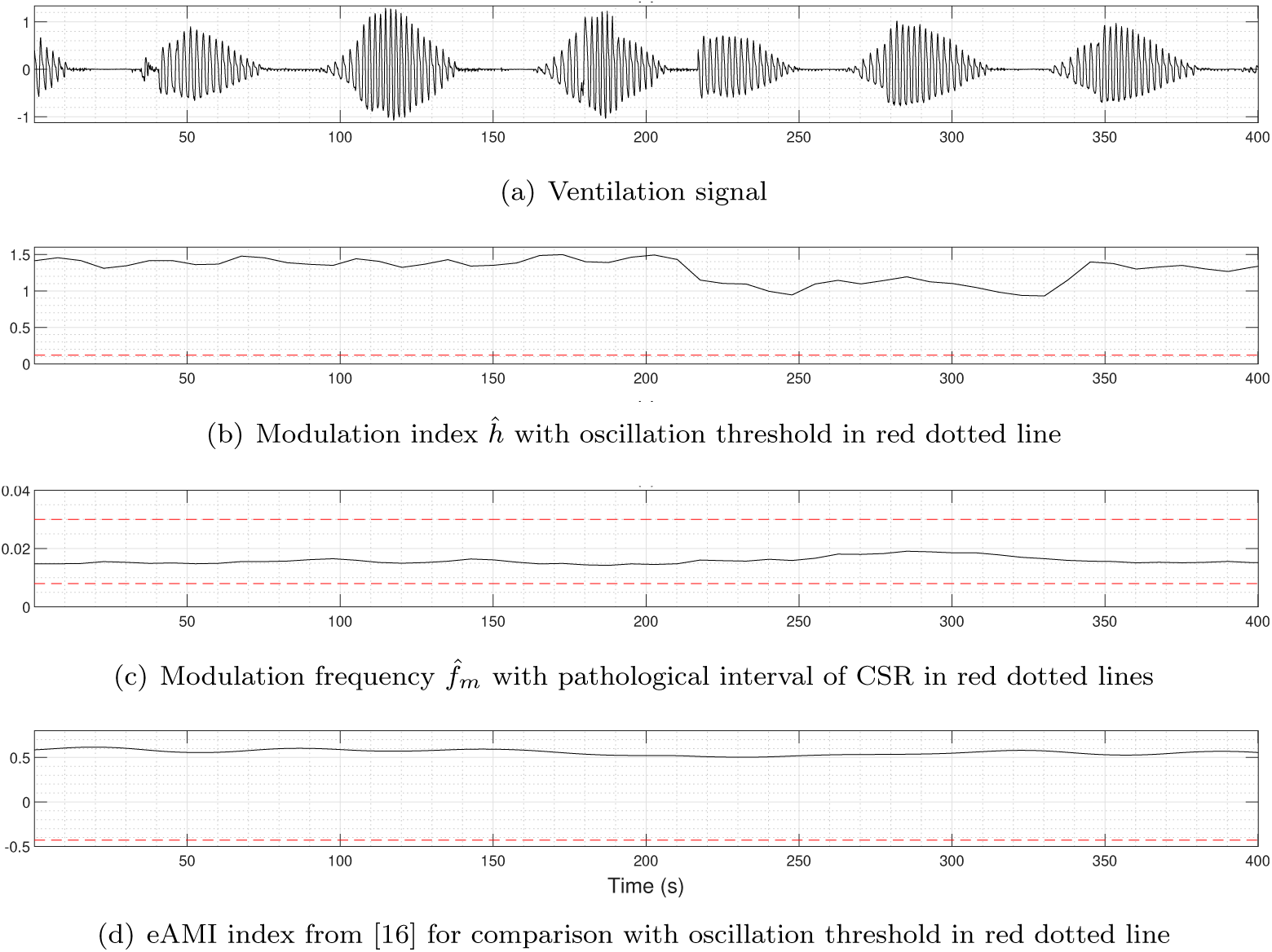
Patient with severe Cheyne-Stokes respiration.

**Fig 5.**
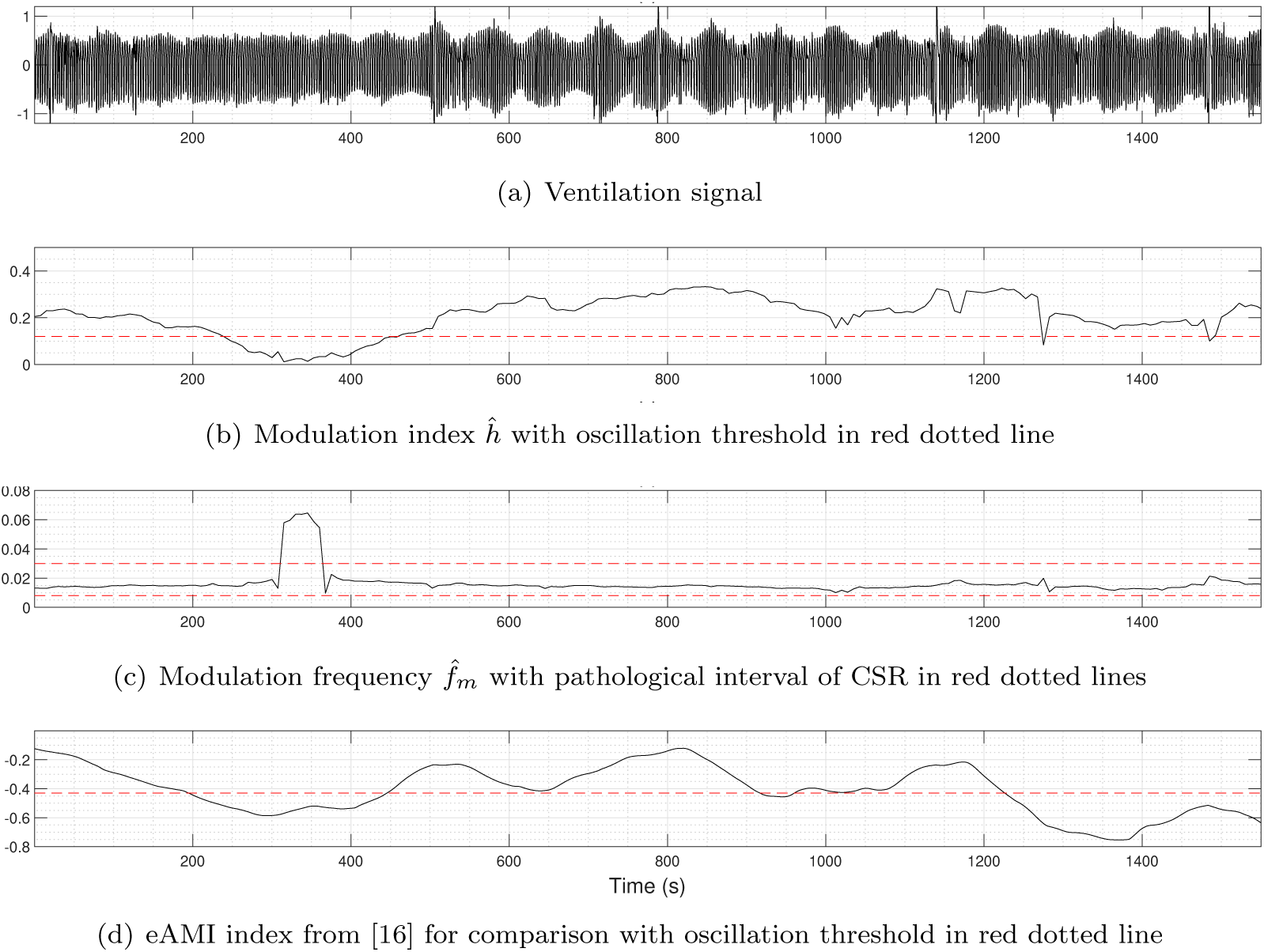
Patient with periodic breathing preceding Cheyne-Stokes respiration.

### 4.5 Confusion matrix

The classification outcomes are described by the confusion matrices presented in Table 2. Each row represents the instance of a predicted class by one of the two algorithms while each column represents the actual class given by experts. If an algorithm performs perfectly with experts, only the diagonal of the matrix will have non-zero values; otherwise, non-zero values outside of the diagonal will specify the class involved in the misclassification. Patients with CSR-CSA are clearly well detected. Patient with diagnosed PB (periodic breathing) is correctly detected so as patients with non-CSR respiration. The interesting fact is that our algorithm classified the undiagnosed three patients in PB.

**Table 2.** Comparison of confusion matrices of our method and the eAMI algorithm [16] on a group of 15 patients.

| Our algorithm | Experts |  |  | eAMI | Experts |  |  |
| --- | --- | --- | --- | --- | --- | --- | --- |
|  | CSR | PB | Non-CSR |  | CSR | PB | Non-CSR |
| CSR | 4 | 0 | 0 | CSR | 4 | 0 | 0 |
| PB | 0 | 1 | 3 | PB | 0 | 1 | 7 |
| Non-CSR | 0 | 0 | 7 | Non-CSR | 0 | 0 | 3 |

CSR: Cheyne-Stokes respiration.
PB: Periodic Breathing typically preceding CSR.

## 5 Discussion

The prevalence of sleep-disordered breathing (SDB) in adults is important and the diagnosis can be challenging as symptoms can be confused with or masked by other pathologies typically associated with SDB. Concerning patients with severe heart failure, CSR has been proven to be a strong factor of higher mortality, thus an early detection is crucial. Periodic breathing is considered to be the early pattern of CSR with small and subtle manifestations tough to detect.

In the present study, the proposed detection method of CSR and PB patterns has shown reliable results by our amplitude demodulation technique applied to patients with severe heart failure. Modeling the ventilation envelope with amplitude modulation leads to characterize its morphological aspect and provides an efficient numerical biomarker of CSR severity. Combining the value of the modulation index *h* and the modulation frequency *f*_*m*_ allows to precisely describe the signal. If *h* is above the pathological threshold experimentally set at *h*_0_ = 0.12 and if *f*_*m*_ shows correlation by being contained in the interval [8, 30] mHz for at least 10 minutes per hour, then a pathological modulation is sufficiently present to be marked as CSR patterns. However, it is important to combine both parameters together, if *h* is above *h*_0_ but *f*_*m*_ is out of the interval or is not stable within, there is no oscillation zone marked. On the contrary, if *f*_*m*_ is contained within the interval but *h* is less than *h*_0_, there is no readable modulation in the envelope. The right way to analyze the parameters is to read carefully the values of *h* first. If it is steadily above *h*_0_ then a modulation is present but we cannot conclude about its nature; if it is under *h*_0_, there is no modulation amplitude in the signal. Then, read *f*_*m*_ signal: if it is continuously contained in the pathological interval, then an oscillation zone is detected; if *f*_*m*_ is unstable (both within and outside the interval for no more than one minute), no modulation is detected. Of course, if it is clearly outside the interval, no modulation is detected.

Finally, if a CSR pattern is detected, the value of *h* is an indicator of severity of the pathology. If 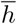, the mean value of *h* during oscillation zones, is in the interval [*h*_0_, 1] then a periodic breathing is present without apnea. The closer 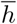 is to 1, the more acute is the pathology. If 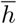 is above 1, then the patient presents a CSR pattern with apnea. The higher 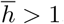, the longer are the apneas and the more severe is the pathology.

Our method achieves better overall results than eAMI. Our final classification accurately detected all patients presenting CSR patterns with or without apnea. It also correctly classified non-CSR patients. Three patients were classified by the expert as non-CSR but with possible early CSR patterns. Those three patients were classified by the algorithm as periodic breathing preceding CSR. The algorithm highlighted the same patients as the experts and allowed to clearly quantify and qualify their breathing to confirm the suspicion of the experts.

The proposed method can be used to monitor periodic breathing through night to determine its progression according to sleep stages or through different exams to observe the evolution within months. It can be a powerful tracker to locate the patient on the continuum of the pathology and help the expert to precisely estimate the evolution of the patient’s symptoms. Also, as our index can be considered as a continuous signal, precising the severity of the modulation through night, it has the advantage over the AHI index that describes the whole process instead of computing the sum of events. It is also an automatic method that does not require any human intervention contrary to AHI estimation.

Finally, the algorithm is based on the same tools that the expert uses: morphology using *h* index that matches the crescendo-decrescendo pattern of periodic breathing and temporal intervals with *f*_*m*_ that specifies the exact frequency of the oscillation.

## 6 Conclusion

We presented a new computational method to detect early patterns of Cheyne-Stokes respiration and to estimate severity levels of pathology from ventilation signals measured on patients. All the components of the proposed method have been tested on a panel of 15 patients. The change point analysis technique has proved to be efficient to detect breathing cycles and the matrix pencil method has provided accurate estimation of the CSR model parameters. Two of them were used to detect CSR zones and to classify the seriousness of the pathology. The classification results showed promising performances of the proposed solution and demonstrated the proof of concept since all the predictions are consistent with experts’ conclusions. A short-term perspective will focus on the possibility to adapt our method to be applied directly on electrocardiogram signals. The mid-term goal is to carry out a clinical study to analyze the cost-efficiency, validate the proposed solution in a larger panel of patients, and propose a robust tuned threshold for the detection.

## Footnotes

1 Left Ventricule Ejection Fraction.

